# A microcontroller-based approach to control oxygen and establish flexible acclimation regimes

**DOI:** 10.1101/2024.06.03.597140

**Authors:** Stefan Mucha

**Affiliations:** Behavioral Physiology, Institute of Biology, Humboldt-Universität zu Berlin, 10099 Berlin, Germany

## Abstract

Environmental control systems are important tools for experimental researchers studying animal-environment interactions. Commercial systems for measurement and regulation of environmental oxygen conditions are relatively expensive and can not always be adapted to varying experimental applications. Here, we present a low-cost and highly flexible oxygen control system using Arduino microcontrollers in combination with a commercial optical oxygen sensor. We show dissolved oxygen (DO) measurements from three experiments that were conducted with mormyrid fish with different DO regimes: 1) acute challenge with defined rates of DO change from normoxia (> 90% air saturation) to hypoxia (10% air saturation), 2) acute stepwise hypoxia in a shuttle-box setup, and 3) long-term hypoxia (15 % air saturation) that was maintained for eight weeks in the home tank of the fish. Overall, the microcontroller-based DO control systems produced the desired experimental conditions with a high and inter-trial correlation (Pearson’s R ≥ 0.987) and sufficient precision, except for the shuttle-box setup, where the lowest DO step of 10% air saturation could not be reached.

## Introduction

Monitoring and regulation of environmental conditions are at the core of experimental studies investigating animal-environment interactions. An important environmental variable is oxygen availability. Almost all eukaryotes depend on oxygen for their survival (Fenchel 2012). In most terrestrial habitats, oxygen is abundantly available (but see, e.g., the naked mole rat or high-altitude habitats, Monge and León-Velarde 1991; Pamenter 2022). In aquatic habitats, dissolved oxygen (DO) can fluctuate significantly on a small spatial and temporal scale (Graham 1990; Kramer 1984). In addition to naturally occurring conditions of low dissolved oxygen (hypoxia), global warming and environmental degradation (e.g., eutrophication) are projected to lead to an increased frequency and intensity of hypoxic events (Breitburg et al. 2018; Diaz and Rosenberg 2008; McBryan et al. 2013). Many experimental studies test how animals respond to changes in environmental oxygen by exposing them to acute hypoxic stress. Such experiments are often combined with chronic oxygen treatments to control for longer-term effects of DO environment (Ackerly et al. 2018; Ackerly et al. 2023; Borowiec et al. 2015; Borowiec et al. 2018; Borowiec et al. 2020; Collins et al. 2015; Cook et al. 2013; Ding et al. 2013; Lomholt and Johansen 1979; Love and Rees 2002; Pan et al. 2017; Prosser et al. 1957; Reardon and Chapman 2010; Shang et al. 2006).

To establish short-term and long-term DO regimes, automated measurement and control systems have many benefits over manual manipulation of environmental conditions: they create reproducible and stable conditions, they operate continuously over extended periods, and they save time and labor for researchers.

Establishing automated DO control in a laboratory is not trivial. The efficiency of DO control depends on many variables that should be taken into consideration, such as water volume, gas pressure and mode of delivery, surface area for exchange with atmospheric oxygen, temperature, water flow and mixing, and desired DO concentration. And due to weathering of components, no automated control system can work without regular maintenance and control of an experimenter. Various manufacturers offer off-the-shelf environmental control systems. While these systems have the benefit of professional support and service, they can easily cost up to 5-6 figures and, once established, are not always adaptable for a different desired experimental design (e.g., establishing constant conditions vs. fluctuating DO, setting stable conditions vs. defined change rates, interfacing with a PC vs. standalone logging and control). The inaccessibility of DO control systems limits ecophysiological research, as it allows only few, well-funded laboratories to conduct studies with controlled DO regimes. Just recently, this issue was addressed by Ern and Jutfelt, who published an inexpensive do-it-yourself system for regulating water oxygen (Ern and Jutfelt 2024). Here, we present an alternative framework for automated DO measurement and control using Arduino microcontrollers as control units (https://www.arduino.cc/) and a commonly used optical DO meter as sensory unit (FireStingO_2_, PyroScience GmbH, Aachen, Germany). We present three different hardware configurations, using solenoid valves, mass-flow controllers and adjustable flow-through valves, and show data from acute hypoxia challenge protocols (steady-rate and stepwise DO decline) and long-term hypoxia treatments (8-week hypoxia acclimation).

## Methods and Results

### General Methods

The hardware backbone for the control system consists of an Arduino-type microcontroller, a FireStingO_2_ sensor, and a gas valve that can be operated by the microcontroller. The serial connection between microcontroller and FireStingO_2_ is established simply by connecting the ‘receive’ pin (RX) of the microcontroller with the ‘transfer’ pin (TXD, pin 4 on connector X1, Fig. 1), and the TX pin of the microcontroller with the RXD pin (pin 5) of the FireStingO_2_. Importantly, microcontroller and FireStingO_2_ have to operate on the same electrical ground for the serial communication to work. This can be easily achieved by powering the FireStingO_2_ directly through the microcontroller 5V-output. To connect the devices, generic jumper wires and a 7-pin PCB connector (Phoenix Contact PTSM 0,5/ 7-P-2,5, Art. No. 1778887) are sufficient. Specific components and additional peripherals can vary depending on the application (table 1). In general, DO control follows a classic cybernetic approach: i) the microcontroller sends commands over the serial connection to measure and read out DO values from the sensor, ii) received DO values are compared to a set value, iii) a control output is calculated (e.g., using proportional-integral-differential control algorithms), and valves are operated to regulate gas flow into the water, iv) steps i-iii are repeated in a defined interval. To streamline the communication between microcontroller and oxygen sensor, we created a small library that contains functions for measurement and readout of DO values (https://github.com/muchaste/ArdOxy). Other functions that were used to record/display measurements, compute control outputs (all control outputs were computed using the PID library by Brett Beauregard, https://github.com/br3ttb/Arduino-PID-Library), and operate gas valves are not integrated in the library but are incorporated into the example Arduino control programs (sketches) that accompany the ArdOxy library. The Arduino sketches that were used in the experiments presented here can be downloaded from figshare (https://doi.org/10.6084/m9.figshare.25764879.v1). Currently, calibration of the oxygen sensor is not included in the library and has to be done using Pyro Workbench (PyroScience) and following the manufacturer’s instructions.

**Figure 1.**
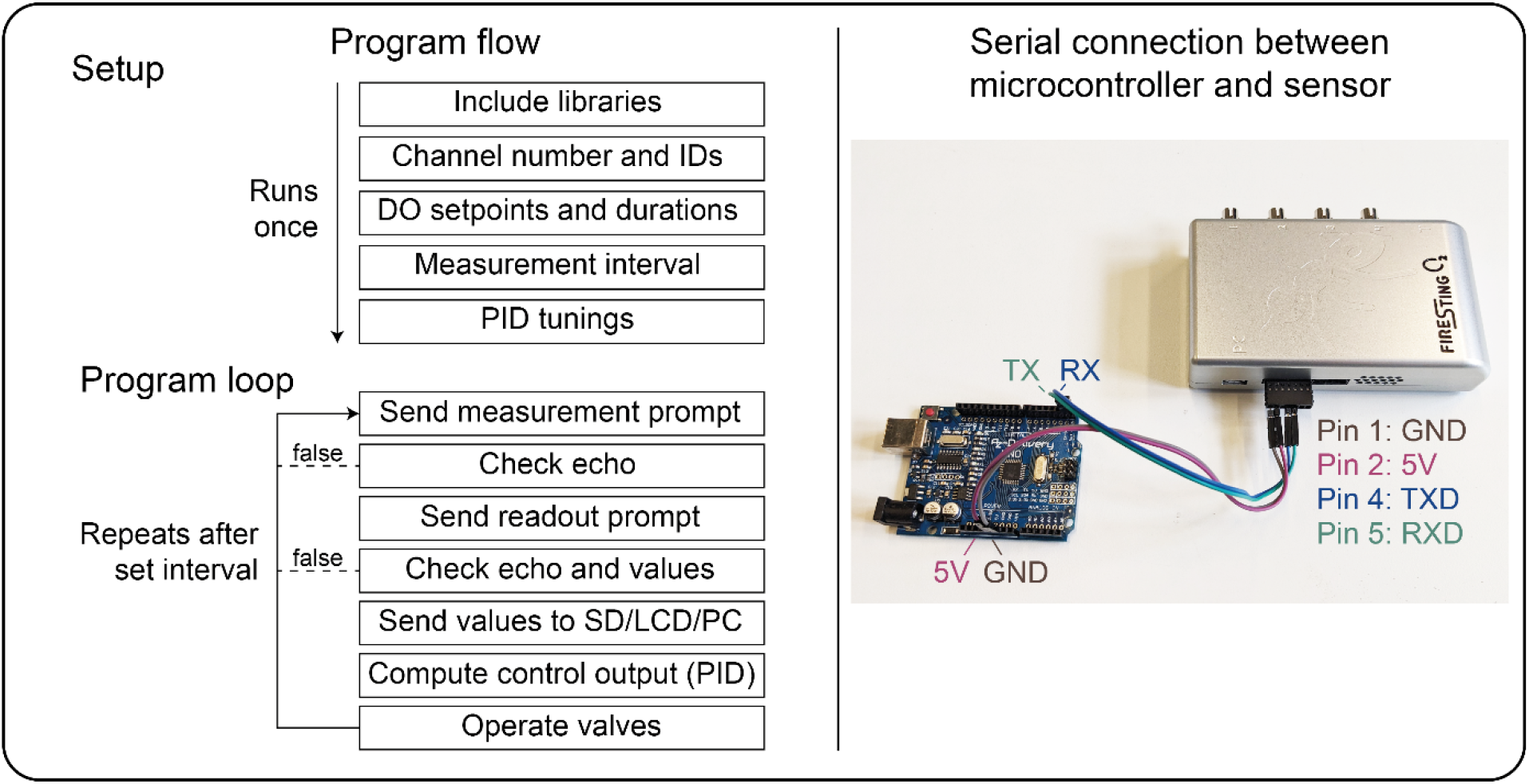
Left: Typical program flow of the control system. First a setup routine is executed once, followed by a program loop that is repeated after a defined interval. To check for errors that can occur in the serial communication between the microcontroller and the sensor, responses of the sensor (echoes) are compared with sent prompts and if there is a mismatch, the program loop can be interrupted or restarted. Right: wiring example for connecting an Arduino Uno-type microcontroller to the FireStingO_2_ sensor via serial connection. Note that the receive-pin (RX) of the microcontroller is connected to the transfer-pin (TXD) of the sensor and vice versa. DO: dissolved oxygen, GND: ground, PID: proportional-integral-differential controller, RX/RXD: receive-pin, TX/TXD: transfer-pin.

**Table 1.** Components of the oxygen control systems. Prices are estimated based on a survey of online shops (as of May 2024).

| Experiment | Component | Model | Price estimate<br>excl. O<sub>2</sub> sensor<br>(USD) |
| --- | --- | --- | --- |
| 1. Acute hypoxia<br>challenge with<br>defined change<br>rates | Microcontroller | Arduino Uno R3 | 25 |
|  | Motor shield | Adafruit Motor Shield V2 | 20 |
|  | Stepper motor | NEMA-17 12V 350mA | 15 |
|  | Shaft coupling | Generic | 5 |
|  | Flow regulator valve | Shuny 6mm push-in valve | 10 |
|  | 12V power supply | Voltcraft VC-11320935 | 15 |
|  | Cables and supplies | Generic | 20 |
|  | **Total** |  | **110** |
| 2. Stepwise hypoxia<br>in shuttle-box | Microcontroller | Arduino Uno R3 | 25 |
|  | Mass-flow controller | MKS 1259-V-10K-S | 450 |
|  | 15V Power supply | XP Power ECL30UD02-S | 72 |
|  | Cables and supplies | Generic | 20 |
|  | **Total** |  | **567** |
| 3. Long-term<br>hypoxia | Microcontroller | Arduino Mega 2560 | 50 |
|  | Display | Adafruit LCD Shield kit | 25 |
|  | SD card logging | Adafruit Data Logger Shield | 15 |
|  | 4 relay board | BerryBase RELM-4 5V | 5 |
|  | 4 solenoid valves | Bürkert 6011 24V (art. 163521) | 240 |
|  | 5V power supply | Voltcraft VC-11258705 | 20 |
|  | 9V power supply | Voltcraft VC-11258705 | 20 |
|  | Housing | Bettermann 2007125K | 30 |
|  | Cables and supplies | Generic | 20 |
|  | **Total** |  | **425** |

We present data from three experiments that were conducted with mormyrid fish as application examples for the control system. A full list of components and manufacturers can be found in table 1. R version 4.0.4 (https://www.r-project.org/) was used for data analysis. Figures were formatted with Adobe Illustrator (Adobe Inc., San Jose, USA). All Arduino sketches and R scripts can be downloaded from figshare (https://doi.org/10.6084/m9.figshare.25764879.v1). All experimental procedures involving animals were approved by the Landesamt für Gesundheit und Soziales Berlin, Germany (Reg. 0278/17).

### Application 1: Acute Hypoxia Challenge in Aquarium with Fixed DO Change Rates

In this experiment, we changed DO in the home tank of the fish (80x30x35cm, water volume of 70L) from normoxic values (>90% air saturation) to 10% air saturation and back to 95% air saturation at a defined change rate. During the trials, water in the tank was mixed with a small pump (CompactON300, Eheim, Deizisau, Germany) and bubble wrap was placed on the water surface to reduce the diffusion of atmospheric oxygen into the water. The DO control system consisted of an Arduino Uno R3 that was connected to a FireStingO_2_ sensor via serial connection (Fig. 1, 2A). A temperature probe (TSUB-21, PyroScience) and a robust DO probe (OXROB 10-CL4, PyroScience) were placed approximately in the middle of the tank. Temperature and DO were measured every two seconds, values were sent from the Arduino to a PC via USB connection and visualized using SerialPlot (https://github.com/hyOzd/serialplot, Fig. 3A). The gas flow was regulated through a needle valve that was connected to a stepper motor via a shaft coupler. A motor shield (Motorshield V2, Adafruit, New York, USA) was used to drive the motor via the Arduino. DO was decreased from normoxia (>95% air saturation) to 10% air saturation within 120 min and maintained at 10% for 100 min by bubbling in nitrogen. Afterwards, DO was increased to 95% within 120 min by bubbling in air. Data from a total of seven trials is shown here (average temperature: 24.9 ± 0.5 °C).

**Figure 2.**
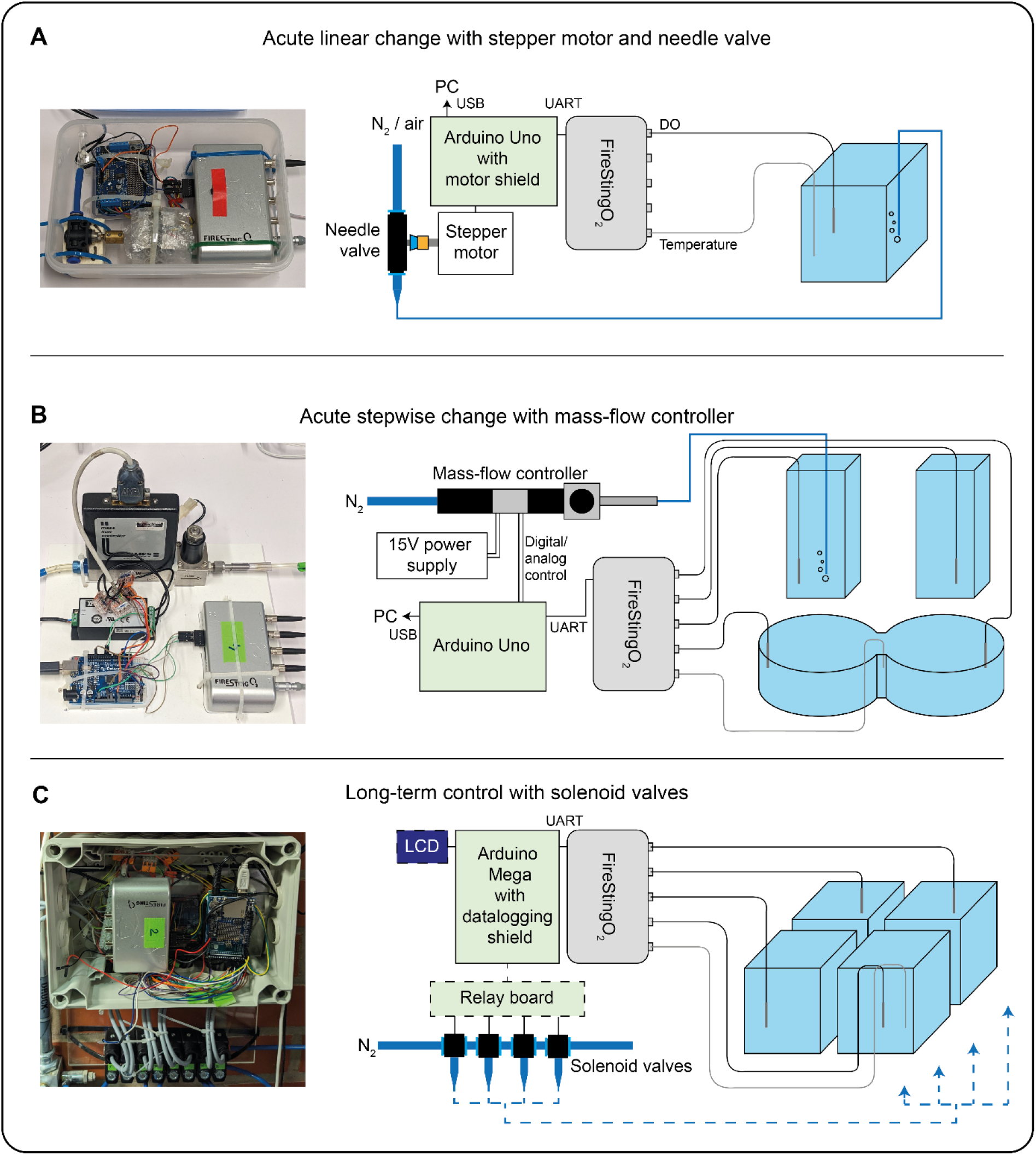
Pictures (left) and schematic representations (right) of three different hardware configurations for DO control. C: the relay board and LCD of the long-term control system are not visible in the picture.

**Figure 3.**
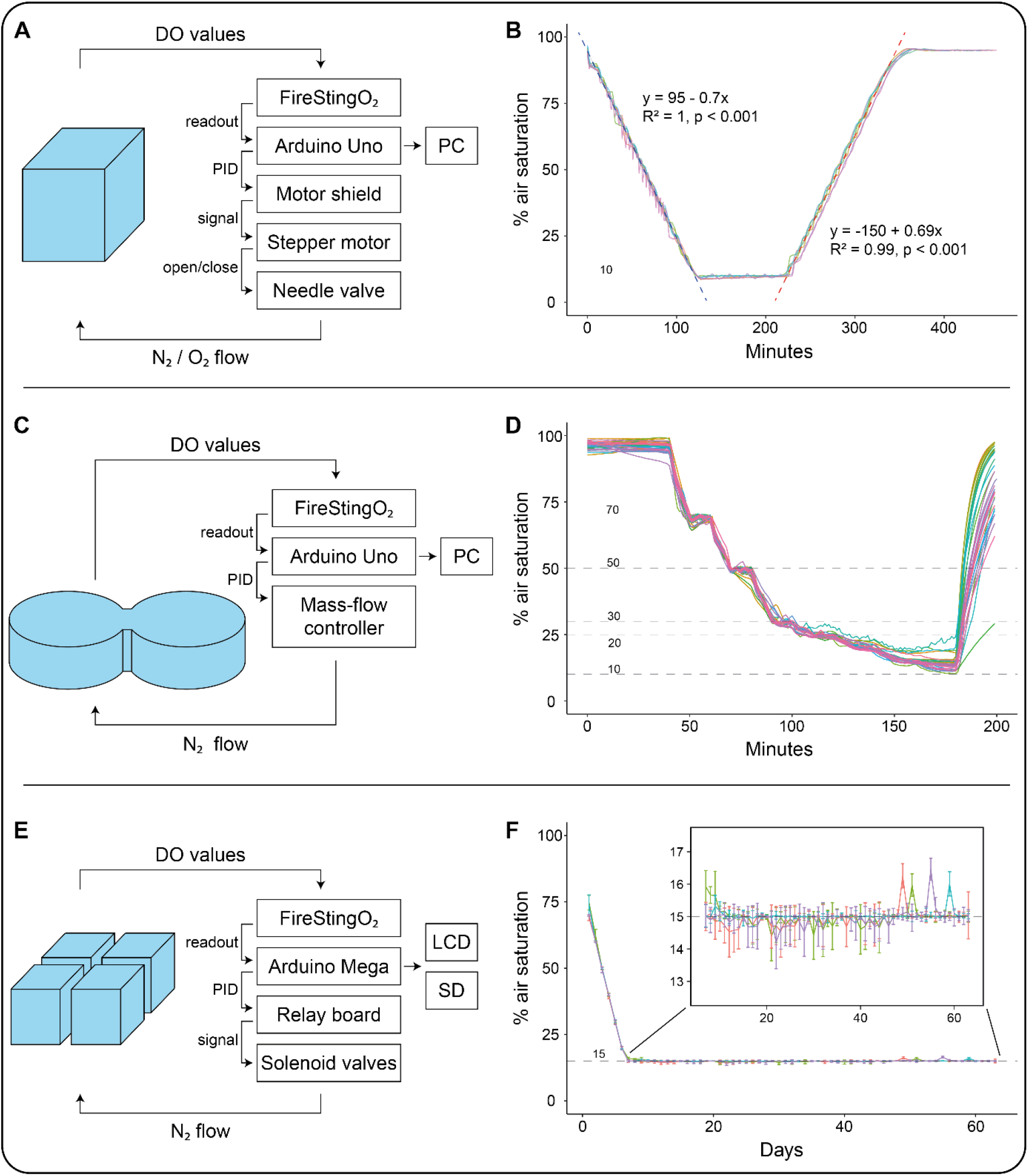
Schematic representations of control cycles (left) and DO measurements (right) of the three experiments. A-B: Acute change with fixed rates in the home tank of the fish. Pearson’s correlation coefficients are shown for the linear change periods of the trial. C-D: Acute stepwise change in a shuttle-box. E-F: Long-term hypoxia in the home tanks of the fish (daily means, error bars show standard deviation). Dashed horizontal lines show target DO setpoints. PID: proportional-integral-differential controller.

Desired values of 10% and 95% air saturation were reached within 120 minutes and DO trajectories were highly correlated across trials (Pearson’s R >= 0.999, Fig. 3B). However, DO curves did show a step-pattern with partial overshoots, especially during DO reduction. During maintenance of 10% air saturation the PID control produced slightly more hypoxic conditions than desired (9.67 ± 0.47% air saturation), normoxic values were reached with higher accuracy (95.06 ± 0.19% air saturation).

Overall, this experiment is an example for a relatively simple application of DO control. The water surface-volume ratio was small, which reduced undesired influx of atmospheric oxygen. The tank was not compartmentalized, which was beneficial for water mixing and resulted in low latencies between adjustment of gas flow and changes of DO. Thus, the system could measure DO and adjust gas flow with a relatively high sample rate of 0.5 Hz, which improves the performance of the PID control algorithm. The motorized needle valve created a more continuous gas flow than e.g., when using a solenoid valve. This is beneficial when the fish might be disturbed by sudden intervals of nitrogen bubbling. We used a shaft coupling to connect a stepper motor to a manual needle valve and thus created a cheap motorized valve which worked reliably after all components and the rotational position of the motor shaft were fixed. The magnitude of overshoots during change of DO was reduced by adjusting the tunings of the PID control algorithm and reducing nitrogen pressure at the outlet.

### Application 2: Acute Hypoxia in a Shuttle-Box with Stepwise DO Change

In this experiment, DO was decreased in a stepwise sequence in a shuttle-box tank (Loligo Systems, Viborg, Denmark) that was adapted from Mucha et al. (2021). The area of the shuttle-box where the fish was located consisted of two round compartments with a diameter of 50cm each, that were connected through a passage (width: 7.5cm, length: 10cm). In each compartment, water was exchanged with a pump (Universalpumpe 1046, Eheim) between the compartment and a buffer tank in which a gas diffusor was placed to bubble nitrogen into the water. A total water volume of 60L was used. Plexiglass lids were placed on the water surface in the compartments of the shuttle-box to reduce the diffusion of atmospheric oxygen into the water. The DO control system consisted of an Arduino Uno R3 that was connected to a FireStingO_2_ sensor via serial connection (Fig. 1, 2B). Per side of the shuttle-box, one robust DO probe was placed in the round compartment and one robust DO probe was placed in the corresponding buffer tank. The temperature probe was placed in the passage between the compartments of the shuttle-box. In the buffer tanks, DO responded with a short latency to gas input. However, because the water exchange rate between the buffer tanks and the round compartments was limited, the latency between gas input and change of DO in the compartment was high, and DO values varied substantially between the buffer tanks and the compartments. As a trade-off between accuracy and latency, we calculated a 1:2 weighted average of DO measured in the buffer tank and DO measured in the compartment. The weighted average was then compared to the set value and gas flow was adjusted accordingly. Temperature and DO were measured every ten seconds, values were sent from the Arduino to a PC via USB connection and visualized using SerialPlot (Fig. 3C). The gas flow was regulated through a mass-flow-controller (1259-V-10K-S, MKS Instruments Deutschland GmbH, München, Germany) that was powered by a +/-15V power supply and controlled via the analog output of the Arduino. Air saturation was lowered in one side of the shuttle-box to 70, 50, 30, 25, 20, 15 and 10% while normoxic conditions were maintained in the other side. Each air saturation value was maintained for 10 min and then changed to the next value within 10 min. Data from a total of 32 trials are shown here (average temperature: 23.4 ± 0.6°C).

DO values showed much higher variability than in experiment 1, although they were still highly correlated across trials (Pearson’s R >= 0.987, Fig. 3D). The reduction to 15% air saturation was successful in 27 of 32 trials but 10% air saturation was not reached (14.13 ± 2.63% air saturation). Overall, DO control in the shuttle-box was much more challenging than in a simple tank. This was mainly due to three factors: large surface-volume ratio, poor water mixing between compartment and buffer tank, and undesired water mixing between hypoxic and normoxic compartments of the shuttle-box. We minimized influx of atmospheric oxygen by covering open water surfaces with lids. Further, we elevated the shuttle-box relative to the buffer tanks to raise the water level (i.e. the proportion of total water volume) in the buffer tanks where the surface-volume ratio was lower and DO could be adjusted more directly. Finally, we conducted regular maintenance of all components to ensure symmetrical flow rates in both compartments in order to reduce water exchange between the compartments. One factor that we could not control for was mixing of the water between the compartments due to swimming activity of the fish. Indeed, the trials with highest deviation of measured DO from set values were trials with very active fish that shuttled between the compartments the most often.

### Application 3: Long-Term Hypoxia in Aquaria

In this experiment, we established long-term hypoxic conditions in the home tanks of the fish. Four tanks of the same size as in experiment 1 were used. Bubble wrap was placed on the water surface to reduce the diffusion of atmospheric oxygen into the water and water was mixed with small pumps (Eheim CompactON300). The DO control system consisted of an Arduino Mega 2560 that was connected to a FireStingO_2_ sensor via serial connection (Fig. 2C). A robust DO probe was placed approximately in the middle of each tank. One temperature probe was placed in a randomly chosen tank. All tanks were in a temperature-controlled room to ensure approximately similar conditions across tanks. Temperature and DO were measured every 60 seconds, values were shown on a display (LCD shield, Adafruit) and logged on a SD card (Datalogger shield, Adafruit, Fig. 3E). The gas flow was regulated through solenoid valves (6011, Bürkert GmbH, Ingelfingen, Deutschland) that were controlled through the digital output pins of the Arduino via a relay module (AZ-Delivery Vertriebs GmbH, Deggendorf, Germany). During every measurement loop, the valves were opened for a duration of 0-15 s to bubble nitrogen into the tanks. DO was decreased from normoxia to 70% air saturation within the first day and then reduced in steps of 10% air saturation per day until 15% air saturation was reached, which was then maintained for eight weeks (average temperature: 25.4 ± 0.4°C).

Long-term control created the desired DO conditions over eight weeks quite reliably (14.96 ± 0.38% air saturation) and DO values were highly correlated across tanks (Pearson’s R >= 0.999). There was some variability in DO values that was due to an inconsistent pressure of nitrogen gas. Overall, the biggest challenge of long-term DO control in the home tanks of the fish was not the physical process of DO adjustment (as was the case in experiment 2), but rather the maintenance of the system.

Ensuring reliable supply of nitrogen gas (without pressure fluctuations), as well as the functioning of all components of the system throughout the experimental duration are obvious minimum requirements for stable long-term control of DO. During the nine weeks of this experiment, we encountered weathering of the DO probes, a malfunctioning solenoid valve, and supply shortage of bottled nitrogen gas. Most of these factors can be anticipated by keeping a contingency of spare components and backup nitrogen gas, as well as conducting regular maintenance.

## General Discussion

### Performance

If other conditions, such as gas pressure and temperature are kept stable, high repeatability can be achieved with microcontroller-based DO control. The PID controller can buffer variability within a certain range (e.g., it adapts to the reduced effect of nitrogen influx at low DO levels and increases the control output), and thus outperforms simple proportional control (i.e., the more absolute change of DO content is necessary, the longer the nitrogen valve is opened). One great advantage of using microcontrollers is their flexibility. They can be used to control virtually any valve (solenoid, MFC, needle valve). If other sensors are incorporated, the control programs can be adapted to control other parameters (e.g., Temperature) as well. The oxygen sensor used here (FireStingO_2_) can measure oxygen in water and air, enabling control of experimental conditions for terrestrial and aquatic environments. The accuracy of DO control depends on PID tunings, gas delivery, and other parameters that are not controlled by the system, such as gas pressure, bubble size and water mixing. Without controlling for these factors, direct comparison with the performance of other DO control systems is of limited significance. Many studies report only the setpoints of their DO regime without presenting the actual conditions that were established (e.g., as mean ± standard deviation DO availability), which reduces the amount of data for comparison. Those that do present DO data show roughly similar accuracy (some examples shown in Table 2).

**Table 2.** List of example studies that have reported long-term DO data (values are mean ± standard deviation except for Reardon and Chapman 2010, where values are reported with standard error, SE).

| Reference | Duration | Air saturation |
| --- | --- | --- |
| Ackerly et al. 2023 | 6 weeks | 33.3 ± 0.2% |
| | 8 days | 31.3 ± 0.3% |
| Ackerly et al. 2018 | 8 weeks | 17.5 ± 3.5% |
| Pan et al. 2017 | 3 weeks | 31 ± 3.5% |
| Reardon and Chapman 2010 | 6 months | 16.9 ± 0.1 (SE)% |
| This study | 8 weeks | 14.96 ± 0.38% |

### Optimization and Limitation

In experimental settings with high latency between control pulse (nitrogen inflow) and effect (DO reduction), achieving stable and precise DO levels can be challenging. Our results suggest several measures to optimize conditions for any DO control system: minimize total water volume and surface-volume ratio; ensure effective water mixing; reduce factors that increase latency of DO changes, such as compartmentalization; use the shortest feasible measurement interval (but beware of oversampling); and conduct preliminary tests to determine optimal gas pressure and PID tunings for the specific application.

A notable disadvantage of our approach compared to other systems is the need for some familiarity with the Arduino programming language to adapt the sketches to the experimental conditions and ensure the system operates correctly (e.g., the internal timer is typically stored as a 32-bit unsigned long variable that rolls over every ~49 days and 17 hours, which can cause chaos in long-term applications). Further, choosing and assembling the hardware are up to the experimenter. While this allows for high flexibility, it also increases the workload.

Currently, our example Arduino sketches are designed to either relay measurements to a connected PC or log them to an SD card. Future enhancements could include functionalities such as uploading data via network to a server.

## Conclusion

Our study demonstrates that microcontrollers are well-suited for controlling DO conditions in both short-term and long-term applications with accuracy comparable to other control systems. Recently, another oxygen control system, OptoReg, was published by Ern and Jutfelt (2024). Both our microcontroller-based system and OptoReg offer relatively inexpensive DIY solutions for establishing DO control in a laboratory, using the FireStingO_2_ sensor as the measurement unit. However, there are some important differences. OptoReg uses the analog output of the FireStingO_2_ to operate components such as solenoid valves via an optocoupler when a setpoint is crossed. It features easy-to-setup hardware and includes a graphical user interface, which is a major benefit because it makes it easy to operate without programming and engineering skills.

Our system uses the serial connection to interface with the sensor, allowing for communication checks between devices but also potentially introducing errors when long cables are used (>5m). The microcontroller-based control system’s primary advantage is its high flexibility. It can be programmed for virtually any DO regime and can control a wide variety of output devices (valves, motors, etc.).

Furthermore, it can function as a stand-alone logging and control unit, which is particularly useful in fish holding facilities or field settings where operating a PC is impractical. However, this flexibility comes at the cost of the time and effort required for experimenters to learn the Arduino programming language, adapt or write control code, and assemble the systems.

Both approaches incorporate open and low-cost technology in research, which is beneficial for many reasons beyond adaptability and affordability. It promotes sharing of code and data, which supports open and reproducible science and encourages researchers to engage with, and understand, the nuts and bolts of their experimental setups.

## Acknowledgments

The author wishes to thank Prof. Rüdiger Krahe for financial support and valuable feedback on the manuscript. Further, I would like to thank Ina Seuffert, Dominik Bekaan and Helen von Drenkmann for providing care for the fish that were used in this study.

## References

Ackerly KL, Krahe R, Sanford CP, Chapman LJ (2018) Effects of hypoxia on swimming and sensing in a weakly electric fish. J Exp Biol 221. 10.1242/jeb.172130

Ackerly KL, Negrete B, Dichiera AM, Esbaugh AJ (2023) Hypoxia acclimation improves mitochondrial efficiency in the aerobic swimming muscle of red drum (Sciaenops ocellatus). Comp Biochem Physiol A Mol Integr Physiol 282:111443. 10.1016/j.cbpa.2023.111443

Borowiec BG, Darcy KL, Gillette DM, Scott GR (2015) Distinct physiological strategies are used to cope with constant hypoxia and intermittent hypoxia in killifish (Fundulus heteroclitus). J Exp Biol 218:1198–1211. 10.1242/jeb.114579

Borowiec BG, McClelland GB, Rees BB, Scott GR (2018) Distinct metabolic adjustments arise from acclimation to constant hypoxia and intermittent hypoxia in estuarine killifish (Fundulus heteroclitus). J Exp Biol 221. 10.1242/jeb.190900

Borowiec BG, Hoffman RD, Hess CD, Galvez F, Scott GR (2020) Interspecific variation in hypoxia tolerance and hypoxia acclimation responses in killifish from the family Fundulidae. J Exp Biol 223. 10.1242/jeb.209692

Breitburg DL, Levin LA, Oschlies A, Grégoire M, Chavez FP, Conley DJ, Garçon V, Gilbert D, Gutiérrez D, Isensee K, Jacinto GS, Limburg KE, Montes I, Naqvi SWA, Pitcher GC, Rabalais NN, Roman MR, Rose KA, Seibel BA, Telszewski M, Yasuhara M, Zhang J (2018) Declining oxygen in the global ocean and coastal waters. Science 359. 10.1126/science.aam7240

Collins GM, Clark TD, Carton AG (2015) Physiological plasticity v. inter-population variability: understanding drivers of hypoxia tolerance in a tropical estuarine fish. Mar Freshw Res. 10.1071/MF15046

Cook DG, Iftikar FI, Baker DW, Hickey AJR, Herbert NA (2013) Low-O2 acclimation shifts the hypoxia avoidance behaviour of snapper (Pagrus auratus) with only subtle changes in aerobic and anaerobic function. J Exp Biol 216:369–378. 10.1242/jeb.073023

Diaz RJ, Rosenberg R (2008) Spreading dead zones and consequences for marine ecosystems. Science 321:926–929. 10.1126/science.1156401

Ding Z, Sun P, Hua X, Bai Y, Shang EHH, Wu RSS, Zuo Y (2013) mRNA expression of select hypoxia-inducible genes and apoptotic control genes in zebrafish exposed to hypoxia during development. Polish Journal of Environmental Studies 22:357–365

Ern R, Jutfelt F (2024) The OptoReg system: a simple and inexpensive solution for regulating water oxygen. Conserv Physiol 12:coae024. 10.1093/conphys/coae024

Fenchel T (2012) Anaerobic Eukaryotes. In: Altenbach AV, Bernhard JM, Seckbach J (eds) Anoxia, vol 21. Springer Netherlands, Dordrecht, pp 3–16

Graham JB (1990) Ecological, evolutionary, and physical factors influencing aquatic animal respiration. Am Zool 30:137–146. 10.1093/icb/30.1.137

Kramer DL (1984) The evolutionary ecology of respiratory mode in fishes: an analysis based on the costs of breathing. In: Balon EK, Zaret TM (eds) Evolutionary ecology of Neotropical freshwater fishes, vol 3. Springer Netherlands, Dordrecht, pp 67–80

Lomholt JP, Johansen K (1979) Hypoxia acclimation in carp - how it affects O_2_ uptake, ventilation, and O_2_ extraction from water. Physiol Zool 52:38–49. 10.1086/physzool.52.1.30159930

Love JW, Rees BB (2002) Seasonal differences in hypoxia tolerance in gulf killifish, Fundulus grandis (Fundulidae). Environ Biol Fish 63:103–115. 10.1023/A:1013834803665

McBryan TL, Anttila K, Healy TM, Schulte PM (2013) Responses to temperature and hypoxia as interacting stressors in fish: implications for adaptation to environmental change. Integr Comp Biol 53:648–659. 10.1093/icb/ict066

Monge C, León-Velarde F (1991) Physiological adaptation to high altitude: oxygen transport in mammals and birds. Physiol Rev 71:1135–1172. 10.1152/physrev.1991.71.4.1135

Mucha S, Chapman LJ, Krahe R (2021) The weakly electric fish, Apteronotus albifrons, actively avoids experimentally induced hypoxia. J Comp Physiol A Neuroethol Sens Neural Behav Physiol. 10.1007/s00359-021-01470-w

Pamenter ME (2022) Adaptations to a hypoxic lifestyle in naked mole-rats. J Exp Biol 225. 10.1242/jeb.196725

Pan YK, Ern R, Morrison PR, Brauner CJ, Esbaugh AJ (2017) Acclimation to prolonged hypoxia alters hemoglobin isoform expression and increases hemoglobin oxygen affinity and aerobic performance in a marine fish. Sci Rep 7:7834. 10.1038/s41598-017-07696-6

Prosser CL, Barr LM, Pinc RD, Lauer CY (1957) Acclimation of goldfish to low concentrations of oxygen. Physiol Zool 30:137–141

Reardon EE, Chapman LJ (2010) Hypoxia and energetics of mouth brooding: is parental care a costly affair? Comp Biochem Physiol A Mol Integr Physiol 156:400–406. 10.1016/j.cbpa.2010.03.007

Shang EHH, Yu RMK, Wu RSS (2006) Hypoxia affects sex differentiation and development, leading to a male-dominated population in zebrafish (Danio rerio). Environ Sci Technol 40:3118–3122. 10.1021/es0522579

